# Distinct prior expectations shape tactile and proprioceptive localization

**DOI:** 10.1101/2025.11.17.688784

**Authors:** Hüseyin O. Elmas, W. Pieter Medendorp, Luke E. Miller

## Abstract

When a mosquito lands on your finger, swatting it away requires your brain to calculate its location in the external space, which depends on the body’s 3D posture. Two competing hypotheses explain how the brain solves this challenge: *the integration hypothesis*, where tactile signals are transformed into spatial coordinates by integrating touch and posture information; and *the cueing hypothesis*, where touch merely cues a location on the body whose position is specified via proprioception. Adjudicating between these hypotheses is nearly impossible without modeling the latent factors underlying somatosensory spatial perception. We fill this gap in the present study. We first formalized each hypothesis from a Bayesian perspective: If touch merely triggers proprioceptive localization (cueing hypothesis), tactile and proprioceptive localization should rely on the same Bayesian computations, with identical prior expectations about the mosquito’s spatial location; If they involve distinct Bayesian computational processes (integration hypothesis), distinct prior expectations may shape tactile and proprioceptive localization. To test these predictions, we had nineteen participants localize either proprioceptive or tactile targets on their fingertips. We then fit their data with several Bayesian models of each hypothesis. Models allowing different prior distributions between modalities provided the best fit for most participants, with 17 out of 19 participants showing significantly different prior distributions across modalities. These provide strong computational evidence that tactile and proprioceptive localization rely on distinct computational mechanisms, a conclusion that has important implications for how we understand these everyday behaviors and their neural mechanisms.

**Author Summary:** When a mosquito lands on your finger and you swat it away, your brain must solve a challenging problem: determining where the mosquito is in space based on where it touched your skin and where your finger is positioned. Scientists have debated how the brain accomplishes this. One hypothesis proposes that the brain transforms touch signals by combining them with information about body posture—an integration process. An alternative hypothesis suggests that touch simply signals which body part was contacted, and the brain then locates that body part with only proprioception—essentially treating touch as only a cue. While these hypotheses make different predictions, distinguishing between them using behavior alone has proven difficult because the underlying computations remain hidden. We addressed this by having participants locate either touches on their fingertips or the fingertips themselves, then used Bayesian computational modeling to reveal the spatial expectations guiding these judgments. Our models showed that tactile and proprioceptive localization rely on distinct spatial expectations, with 17 of 19 participants showing significantly different patterns. These findings provide computational evidence that localizing touch involves transformations beyond simply locating the body, supporting the integration hypothesis and challenging the idea that touch merely cues body location.

## Introduction

Tactile localization is such a common everyday behavior that we hardly give it a second thought. Yet, the computational process underlying it remains a mystery. Picture a mosquito landing on your finger, which you want to swat away. The mosquito’s position is initially mapped in a skin-based reference frame that is implemented somatotopically in S1 [1]. However, simply knowing where the mosquito touches your skin is insufficient to swat it away, since movement shifts the same patch of skin to an entirely different position in space. How does the brain reliably locate tactile stimuli?

One prominent explanation for tactile localization, which we refer to as the *integration hypothesis*, proposes that tactile cues are combined with proprioceptive information about body posture [2]. This process remaps touch from the skin-based (somatotopic) to a spatial (spatiotopic) reference frame, often centered on another body part (e.g., eyes, trunk). There are several lines of evidence in favor of the integration hypothesis. For example, tactile localization judgments are affected by body posture, such as gaze direction [3, 4]. Effects of body posture take several hundred milliseconds to influence a tactile decision [5, 6, 7], indicative of a dynamic process. Even non-spatial tasks, such as tactile temporal order judgments, are affected by task-irrelevant limb posture [8, 9, 10, 11]. At a neural level, activity in frontal and parietal cortices represent tactile stimuli in external spatial coordinates [12, 13, 14, 15, 16]. Neurons in these regions often show multimodal tactile-proprioceptive receptive fields, responding to both tactile input and specific body positions [17, 18, 19, 20].

An alternative proposal, which we refer to as the *cueing hypothesis*, suggests that tactile input only serves to indicate a location on the body part [21, 22, 23, 24]. A recent study supporting this idea combined tactile localization with a temporal-order judgment task [23]. Participants pointed in external space to the first of two rapidly presented tactile stimuli, one on each hand. When participants incorrectly identified which hand was first touched, they tended to mislocalize the initial stimulus to the location of that secondly touched hand— rather than the first touched hand—at the time of the first stimulation. The authors argue that touch merely acts as a cue for a proprioceptive location of the body in space and is never remapped. Thus, according to the cueing hypothesis, localizing a touch on the fingertip is computationally equivalent to localizing the fingertip itself—suggesting that tactile localization operates similarly to proprioceptive localization.

Several studies have compared tactile and proprioceptive localization [25, 26, 27, 28]. For example, Rincon-Gonzalez et al. [26] found that localizing touch on the hand preserved the overall spatial structure of estimation errors in proprioceptive hand localization, but the magnitude of errors was reduced. However, drawing strong theoretical inferences about the similarity of underlying mechanisms from behavioral data alone is challenging, as latent computational variables that distinguish between alternative mechanistic accounts—such as those posited above—are not immediately observable without computational modeling [29]. To this end, computational modeling may offer a promising approach to disentangle these sources of bias by relating observed behavior to underlying latent transformations and variables.

In the present study, we approach tactile and proprioceptive localization from a Bayesian perspective [30, 31]. Bayesian models have proven effective for explaining the origins of perceptual errors and biases [32, 33, 34, 35]. Because sensory input is noisy [36], the brain must make probabilistic inferences, combining uncertain sensory evidence with prior expectations [37]. Bayes’ theorem formalizes this process. In terms of localization, the probability of a particular stimulus location *x* given the sensory evidence (the posterior, *𝒫* (*x*|*e*)), is proportional to the likelihood of observing that evidence (*ℒ* (*e*|*x*)), multiplied by the prior probability of the stimulus (Π(*x*)); or, *P* (*x*|*e*) ∝ *ℒ* (*e*|*x*) × Π(*x*). This approach reduces error by weighting evidence by its reliability, but it can also introduce systematic biases toward prior beliefs. Notably, Bayesian models have successfully explained several somatosensory phenomena, such as skin-based tactile localization [38], the cutaneous rabbit illusion [39, 40], arm posture biases [41], and distorted body representations [42].

The cueing and integration hypotheses make competing predictions about the Bayesian computations involved in tactile and proprioceptive localization. The cueing hypothesis essentially equates tactile and proprioceptive localization, predicting they utilize the same prior expectations. In contrast, the integration hypothesis posits that localizing touch requires additional remapping transformations compared to localizing the body. These additional transformations may introduce different systematic biases. Moreover, the natural statistics encountered during tactile versus proprioceptive localization may differ, potentially leading to distinct prior expectations for each modality.

To directly adjudicate between these hypotheses, we designed a spatial localization experiment where participants either localized tactile targets on the fingers (tactile localization) or their fingers themselves (proprioceptive localization). Bayesian models were fit to localization errors in each task to test whether tactile and proprioceptive localization utilize the same (cueing hypothesis) or distinct priors (integration hypothesis). We found that models with significantly distinct spatial priors for tactile and proprioceptive localization provided better fits for most participants. These findings provide computational evidence against the cueing hypothesis and are in support of the integration hypothesis. In the following, we first provide a brief, but more formal Bayesian description of the two hypotheses.

### Theoretical background

Consider the task of localizing a somatosensory target in a two-dimensional space. We can formalize this localization process as 2D Bayesian inference, where the perceived location (posterior, *𝒫*) results from combining sensory measurements (likelihood, *ℒ* ) with prior expectations (Π):

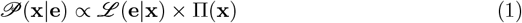

Here **x** denotes the 2D spatial location and **e** the sensory measurements detected by proprioceptors and/or mechanoreceptors. ℒ (**e**|**x**) represents the likelihood function that describes the probability of observing the sensory measurement given target at location **x**, and Π(**x**) represents the expectations over possible target locations, derived from previous observations [43] and internal models [44]. The posterior distribution 𝒫 (**x**|**e**) represents the probability distribution over possible stimulus locations given the sensory evidence; this computation is graphically depicted in Fig 1B.

**Figure 1.**
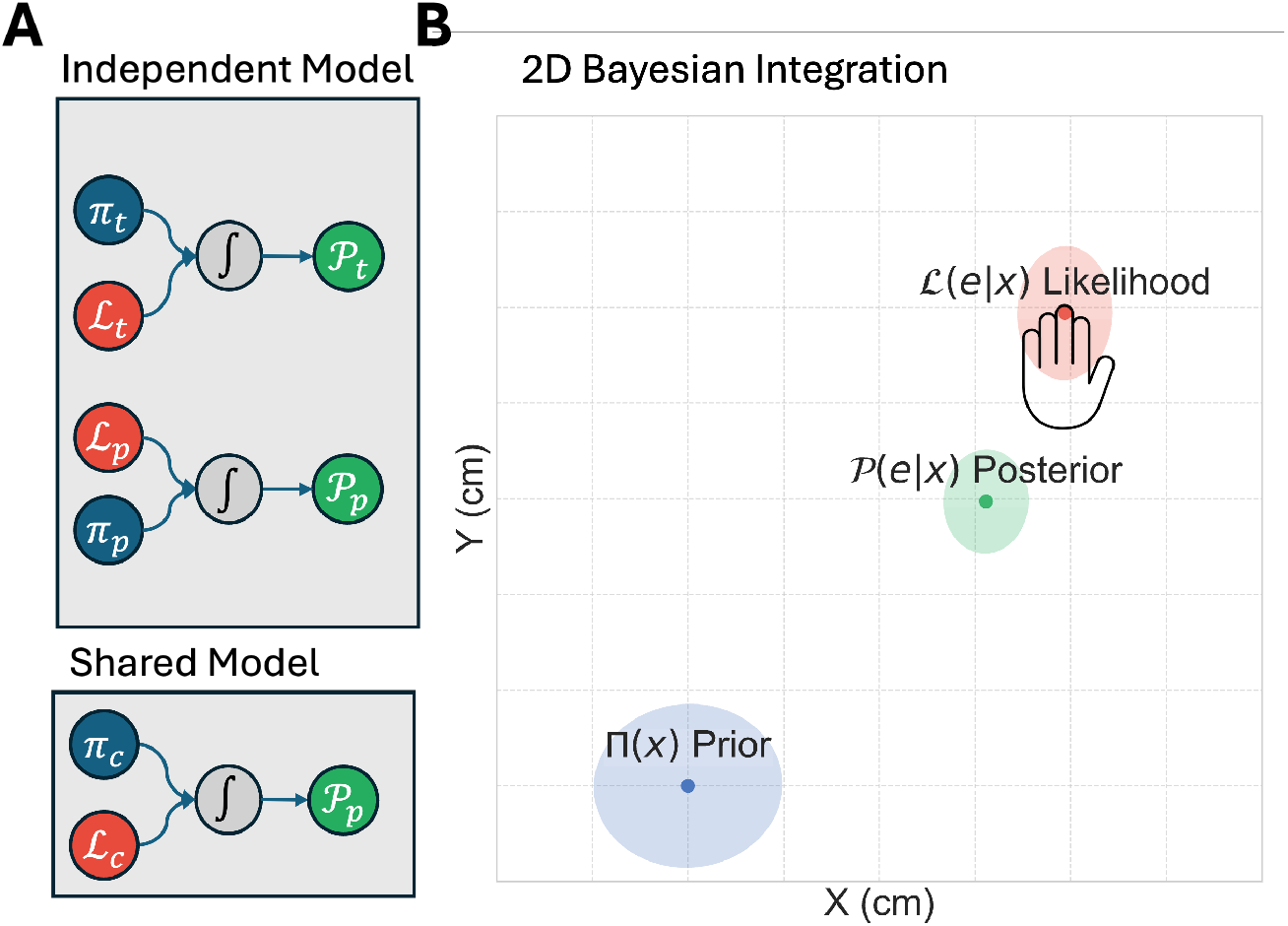
A) Comparison of model variants testing cueing vs. integration hypotheses. The Independent Model (top) allows separate priors (*π*_*t*_, *π*_*p*_) and likelihoods (*L*_*t*_, *L*_*p*_) for tactile (t) and proprioceptive (p) conditions. The Fully Shared Model (bottom) uses identical priors (*π*_*c*_) and likelihoods (*L*_*c*_). The arrows illustrate the computational flow: prior and likelihood are combined via integration (∫) to produce a posterior distribution (*P* ). B) 2D Bayesian inference showing how spatial localization (posterior; green) emerges from combining prior expectations (blue) with sensory evidence (likelihood; red). As can be seen, the presence of a prior leads to a localization error (i.e., offset between likelihood and posterior).

Having introduced the general Bayesian framework, we can now formalize the integration and cueing hypotheses.

### Integration Hypothesis

Consider the task of localizing in space the fingertip or a touch on the fingertip. According to the integration hypothesis, localizing proprioceptive and tactile targets involves distinct computational steps. Proprioceptively localizing the fingertip (**x**_*p*_) requires mapping joint angles (***θ***) and body segment lengths (**l**) to Cartesian coordinates. This can be expressed as:

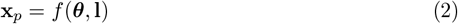

where *f* is a transformation function (e.g. a kinematic chain function) that converts the internal body parameters into external spatial coordinates [41].

Localizing a tactile stimulus on the fingertip **x**_*t*_ requires additional transformations. Here, the mapping involves not only joint angles and segment lengths, but also a function g that maps the stimulus from somatotopic (skin-based) to spatiotopic Cartesian coordinates

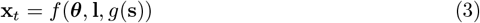

As tactile localization also includes the needed proprioceptive transformations, the additional transformations *g* would likely introduce additional noise and/or biases to the resulting localization. In its strongest form, the integration hypothesis predicts different distinct priors and likelihoods for tactile and proprioceptive localization, i.e., Π (**x**)≠ Π_*p*_(**x**) and likelihoods *L*_*t*_(**e**|**x**)≠ *L*_*p*_(**e**|**x**).

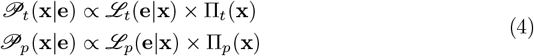

In a weaker version of the integration hypothesis, the tactile and proprioceptive likelihoods share the same uncertainty but still may have distinct spatial priors.

### Cueing hypothesis

In contrast, the cueing hypothesis proposes that tactile and proprioceptive localization rely on the same computational processes:

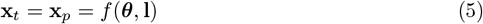

Here, tactile stimulation simply cues a specific body location, which is then localized using the same computational mechanisms underlying proprioceptive localization. Thus, the Bayesian inference underlying localization in both tasks involves common likelihoods and priors:

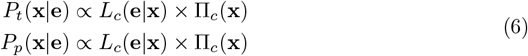

In a weaker version of the cueing hypothesis, tactile and proprioceptive evidence has different levels of noise (e.g., added from the cueing)—and thus distinct likelihoods—while maintaining that both modalities access the same underlying spatial representation and thus share the same spatial prior.

## Methods

### Participants

The experiment initially included 21 healthy adults. Due to technical errors, data from two participants were excluded, leaving 19 participants (ages 17-23, 18 females) in the final analysis. All of them were right-handed, as assessed by the Edinburgh Handedness Inventory (score range: [62.5-100]), reported normal or corrected-to-normal vision, and had no history of neurological disorders. They provided written informed consent before the experiment, which was approved by the Ethics Committee of the Faculty of Social Sciences of the Radboud University. Participants received course credits for their participation.

### Setup

Participants were seated in an adjustable chair, positioned at the edge of a table and facing the short side of a large computer monitor (42” LED iiyama ProLite TF4237MSC-B3AG, Tokyo, Japan, with dimensions 930mm × 523mm, 1920 × 1080 pixels). The computer monitor was placed on its backside in the lengthwise orientation, allowing participants to make judgments in depth. The monitor was securely placed on top of a table with a supportive structure (3 cm from the edge and 24 cm above the table). Participants were positioned with their body midline aligned to the screen center using a reference line displayed at the onset of a block of trials.

The participant’s left hand was placed under the computer monitor and on top of a stimulation platform embedded with three tactile solenoids. The solenoids were arranged to contact with the fingertips of the participant’s index, middle, and ring fingers. The solenoids were placed in a custom-made holder that maintained their stability and upright orientation. This holder was positioned between the monitor and the table surface (∼20cm below the screen). To ensure alignment between the screen coordinates and solenoid positions, we implemented a system using two perpendicular sliding rulers along horizontal and vertical dimensions. These rulers were aligned with the screen and positioned the solenoid holder directly beneath the screen with millimeter precision. The rulers could be locked at any location to prevent movement of the solenoid platform. During the experiment, participants used their right hand to respond with the mouse while their left hand rested on the solenoids.

To eliminate auditory cues from the tactile stimulators, participants wore noise-cancelling headphones (Sony WH-1000XM5) that delivered continuous white noise. Additionally, to eliminate visual feedback of the forelimbs, an opaque cloth covered participants’ hands, forearms, and the stimulation apparatus.

### Experimental Design and Procedure

In separate blocks of trials, participants performed either a tactile or proprioceptive localization task. In each trial of the respective task (Fig 2A), participants received either a tactile target (three brief taps, each lasting approximately 50ms, separated by short intervals, for a total sequence of roughly 0.5 s) or a proprioceptive target, which was indicated by text on the screen specifying one of the fingertips (presentation time: s). Participants indicated the perceived target location on the screen using a mouse pointer, controlled with their right hand. Following target presentation, they had 5 s to respond. If they failed to respond within this period, a ‘Missed’ message appeared briefly on the screen (0.5 s), and the trial was recorded as a miss. Missed trials (∼0.1% of all trials) were excluded from subsequent analyses. No visual feedback about localization accuracy was provided. Trials were separated by a 0.5 s inter-trial interval. During this period, the mouse cursor was invisible. At the start of each trial, the cursor reappeared at a random location within a 5 cm radius circle centered in the lower portion of the screen.

**Figure 2.**
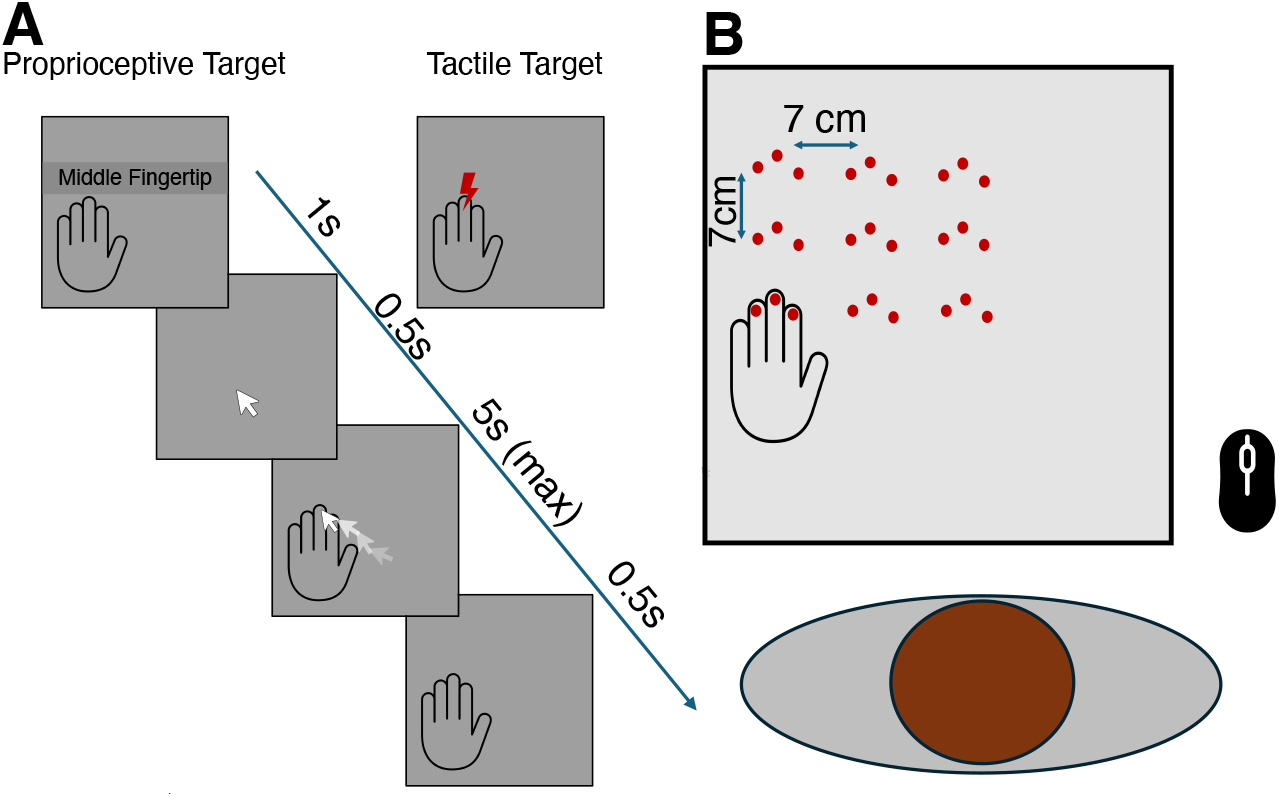
A) Trial structure showing temporal sequence for both tasks. Each trial began with the target presentation (0.5s): tactile stimulation or visual text indicating which fingertip to localize. Participants respond by clicking perceived target location on screen (5s max), followed by 0.5s inter-trial interval. B) Setup with participant positioned at horizontal monitor. The Left hand was placed on a platform beneath the screen with fingertips contacting the solenoids. Red dots show all solenoid positions on the 3×3 grid of hand positions used across blocks.

The experiment comprised 18 blocks of 36 trials, with 9 blocks of tactile localization and 9 blocks of proprioceptive localization, presented in an intermingled order. Hand position varied across a pre-determined 3×3 grid with 9 different locations (Fig 2B), with a spacing of 7 cm between adjacent positions in both horizontal and vertical dimensions; the rightmost column of the grid was aligned approximately with the participant’s midline. In total there were 27 unique target locations (i.e., 3 fingers and 9 hand locations). Block order was randomized for each participant using a random permutation of the 18 blocks. Between blocks, participants could take breaks to prevent fatigue while the experimenter repositioned their left hand to the next location. Each block comprised 36 trials (12 trials per finger), with target fingers randomized within blocks. In total there were 108 trials per finger per task, across different hand locations (324 trials per task; 648 trials in total).

From the perspective of the cueing hypothesis, both tasks involved proprioceptive localization but with different cues: either a tactile cue, or a verbally written cue. In contrast, from the perspective of the integration hypothesis, the participants localized tactile targets and proprioceptive targets within their respective modalities.

### Behavioral analysis

To assess performance within each task, and to descriptively analyze the data, localization judgments on each trial were transformed into an error vector [26, 45]. Error vectors were defined per trial as magnitude and direction of the response location versus the target location in the 2D plane. For each of the nine target locations, and separately per task (tactile, proprioceptive), we calculated the mean ± SD of the magnitude and the circular mean ± circular SD of the direction across trials. For magnitude differences, we performed paired-samples t-tests on target-wise means to assess systematic accuracy differences between modalities. For angular distances, we calculated the absolute circular difference between mean error directions for each participant, then used Watson–Williams tests [46] to assess directional differences. Angular distances range from 0° (identical directions) to 180° (opposite directions). Finally, all trials were pooled across participants to report the grand-mean and SD of error magnitude and circular direction across all target locations.

### Computational Modeling

#### Bayesian Modeling Framework

To quantitatively distinguish between the cueing and integration hypotheses, we modeled participants’ localization responses as the output of a process of Bayesian inference (see Theoretical Background). In our computational modeling, we represented all spatial locations and responses in Cartesian coordinates within the 2D workspace. This modeling approach allowed us to decompose localization behavior into its underlying computational components: sensory uncertainty (likelihood) and spatial expectations (priors), enabling direct comparison of these parameters between tactile and proprioceptive conditions.

Spatial localisation for each task was formalised as 2D Bayesian inference in Cartesian coordinates,

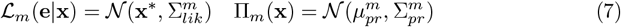

where *m* ∈ { *t, p, c*} denote modality-specific or shared parameters *t* (tactile), *p* (proprioceptive) and *c* (common) to both. **x**^∗^ is a 2D vector, representing the true target location, 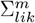 is a 2×2 matrix, capturing the sensory uncertainty and 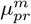 and 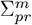 represent likewise the mean and the uncertainty of the prior respectively. Note that we assume that the somatosensory system is well-calibrated to all measurements, and thus the likelihoods are centered on the true location (**x**^∗^ = **x**). For each model variant and modality, the posterior was calculated according to Bayes’ rule:

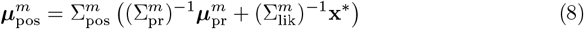

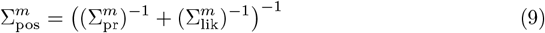

Within the Bayesian modeling framework above, we developed four model variants with different parameter sharing structures to test the competing hypotheses:

*Independent Model* represents the strongest version of the integration hypothesis, where all likelihoods and priors are modality-specific: 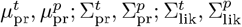

*Shared Likelihood Model* represents a weaker version of the integration hypothesis, constraining likelihood uncertainty, 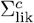, to be identical across modalities while retaining modality-specific priors: 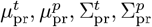

*Fully Shared Model* represents the strongest version of the cueing hypothesis, where the prior and likelihood are fully modality independent, 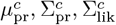

*Shared Prior Model* Represents a weaker version of the cueing hypothesis, imposing a common spatial prior, 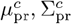, but allowing modality-specific likelihood uncertainties 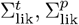

### Model Fitting

To fit our models to the participant responses, we calculated the response distribution, which represents the probability of observing a response given a target location. The response distribution has the same mean with the posterior distribution, and its covariance can be calculated by:

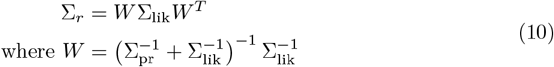

This reflects the fact that behavioral variability stems from the sensory noise and the likelihood’s weight relative to the prior.

We used maximum likelihood estimation to fit each model variant to the individual participant data. For each participant, the log-likelihood was computed as:

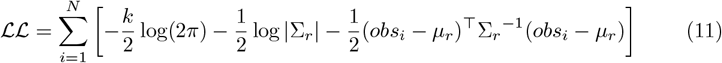

where N is the number of observations, *µ*_*r*_ is the mean and Σ_*r*_ is the covariance of the response distribution, k is the number of dimensions (2) and *obs*_*i*_ is the response location for trial *i*.

To avoid local minima, we used multi-start optimization with 100 random initial parameter sets for each model. Initial parameters were generated using Latin Hypercube Sampling to ensure efficient coverage of the parameter space. Parameter bounds were set to ensure valid covariance matrices and reasonable spatial scales over the workspace. The optimization was done using a custom script using SciPy in Python with L-BFGS-B algorithm [47].

### Model Comparison

To determine which model best accounts for the observed spatial localization behavior, we compared models based on the Bayesian Information Criterion (BIC):

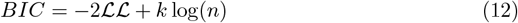

where ℒ ℒ represents Log likelihood value, *k* is the number of free parameters, and *n* is the number of observations.

Next, we computed McFadden’s pseudo-*R*^2^ [48] and Cox-Snell-*R*^2^ [49] to assess model fit quality relative to a null model, where the null model that assumes responses were generated from a single bivariate Gaussian distribution that ignored both target locations and modality differences:

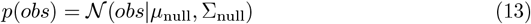

where *µ*_null_ represents the global mean response location and Σ_null_ represents the overall response covariance across all conditions. This model has five free parameters: two for the mean location (*µ*_*x*_, *µ*_*y*_) and three for the symmetric covariance matrix (*σ*_*x*_, *σ*_*y*_, *ρ*). The log-likelihood of this null model was computed as:

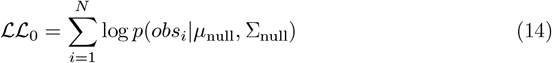

where *obs*_*i*_ represents the response location on trial *i*.

### Comparing Priors

To quantitatively assess differences between tactile and proprioceptive spatial priors, we employed the Wasserstein-2 distance metric [50]. This metric was chosen for its ability to capture differences between probability distributions while maintaining mathematical properties such as symmetry and proper metric behavior. In our case, since we modeled spatial distributions as bivariate Gaussians, we utilized the closed-form solution:

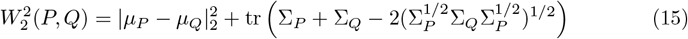

where P and Q are two bi-variate normal distributions, *µ*_*P*_, *µ*_*Q*_ represent the means and Σ_*P*_ and Σ_*Q*_ represent the covariance matrices of these distributions.

To determine statistical significance of observed distances between modality-specific distributions, we implemented a permutation test. For each participant, we computed the true Wasserstein-2 distance between fitted tactile and proprioceptive prior distributions from the Independent Model. We then generated a null distribution by repeatedly shuffling (1000 permutations) modality labels across trials and refitting the model to calculate resulting distances. Model fitting for the permutation analysis employed the same optimization framework as the main analysis (see above), with only 50 random starts per shuffle (instead of 100) to balance computational efficiency. If both modalities share the same underlying spatial representation, randomly reassigning trial labels should produce distances similar to the observed distance. In contrast, a significantly larger observed distance would indicate non-trivially distinct spatial representations for tactile and proprioceptive localization. Statistical significance was assessed by computing the proportion of null distances exceeding the observed distance, with significance threshold set at p < 0.05.

## Results

### Behavioral Results - Descriptive Analysis

We first examined localization performance to assess whether tactile and proprioceptive conditions produced systematically different error patterns. Fig 3 displays the global mean error vectors for proprioceptive (red) and tactile (blue) task across the target locations. As shown, and consistent with prior findings [25, 26, 27, 28], the magnitude and direction of errors differed between modalities at most locations.

**Figure 3.**
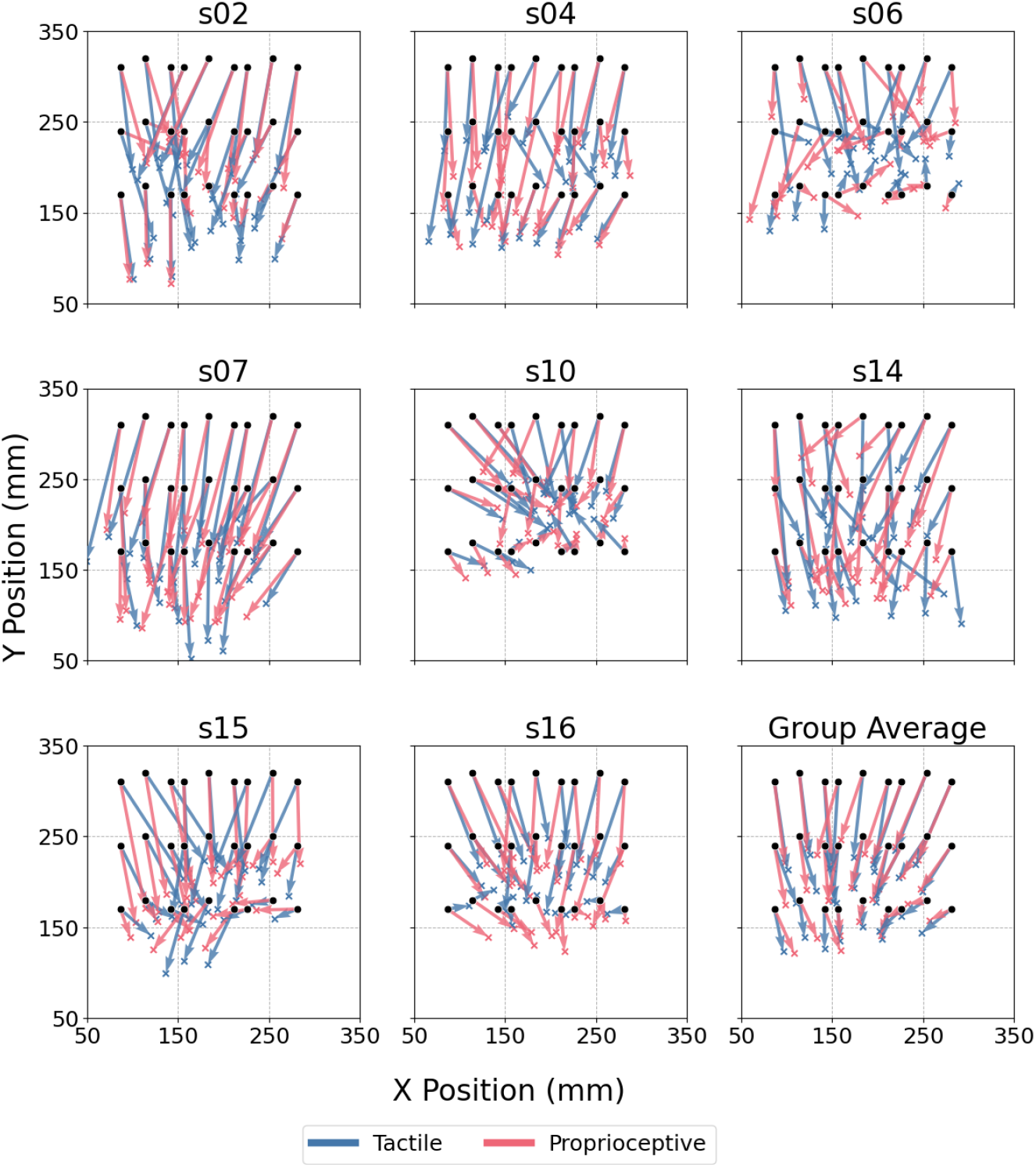
Mean error vectors for proprioceptive (red) and tactile (blue) task over all spatial locations. Error vectors as arrows across a 3×3 grid of target locations. Each arrow originates from a target position (black dots) and points toward the mean response location, with red arrows representing proprioceptive errors and blue arrows representing tactile errors.

Quantitatively, the mean between-modality difference in error magnitude was 2.0 mm (SD: 11.0 mm), with a mean directional difference of 15.0^°^ (SD: 6.0^°^). Individual participant analyses revealed heterogeneous patterns of results: paired t-tests on error magnitudes identified significant modality differences in 8 of 19 participants (42.1%;, p < 0.05), while Watson–Williams tests on error angles showed significant differences in 11 of 19 participants (57.9%;, p < 0.05). While not all participants exhibited statistically detectable differences in both error components, a substantial proportion showed significant modality-specific patterns in either error magnitude or direction (16 of 19 participants (84%), showed at least one effect; Fig 4).

**Figure 4.**
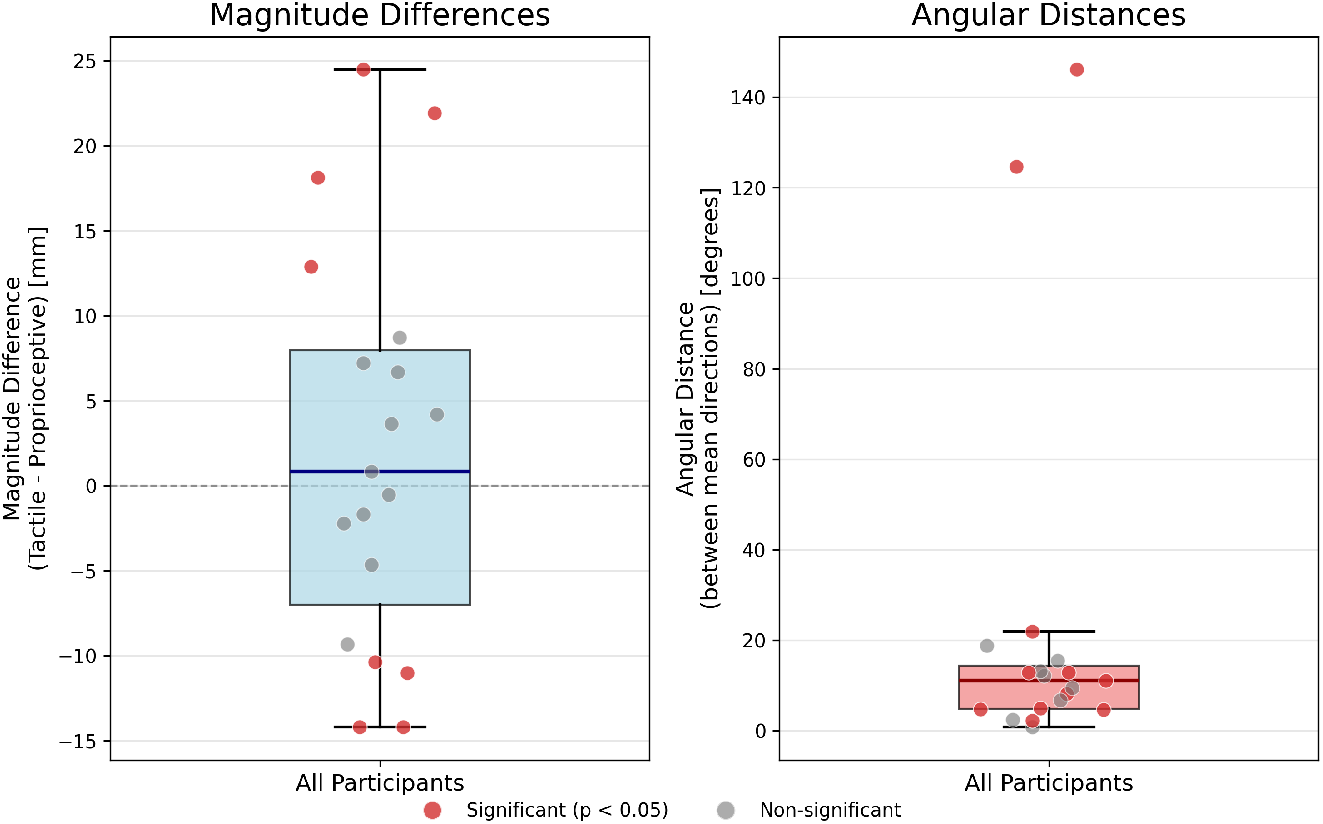
Differences in localization error across participants. Magnitude differences (left) and angular distances (right) between tactile and proprioceptive localization errors for individual participants. Angular distance represents the absolute circular difference between mean tactile and proprioceptive error directions for each participant. Red dots indicate significant differences (p < 0.05), gray dots indicate non-significant differences. Box plots show the distribution across all participants.

These results replicate previously observed differences in the error patterns between tactile and proprioceptive localization [25, 26, 27, 28]. As in these studies, these results alone cannot identify the underlying reasons for these differences. Therefore, to determine the underlying computational basis of these modality-specific error patterns—and to arbitrate between the cueing and integration hypotheses—we applied Bayesian modeling to each participant’s localization behavior.

### Computational Modeling Results

We first evaluated model performance by comparing the predicted and observed response patterns across participants. We systematically compared models representing different theoretical positions, beginning with the Independent and Fully Shared models as extreme versions of the integration and cueing hypotheses. The Independent Model outperformed the Fully Shared Model in 18 of 19 participants (mean ΔBIC: -210 range: [-425, 4]; mean BF: 4.58 × 10^45^; Fig 7A). This is clear in Fig 6 and Fig 5, which compares the model predictions with the observed responses of several participants, for both tactile and proprioceptive localization respectively.

**Figure 5.**
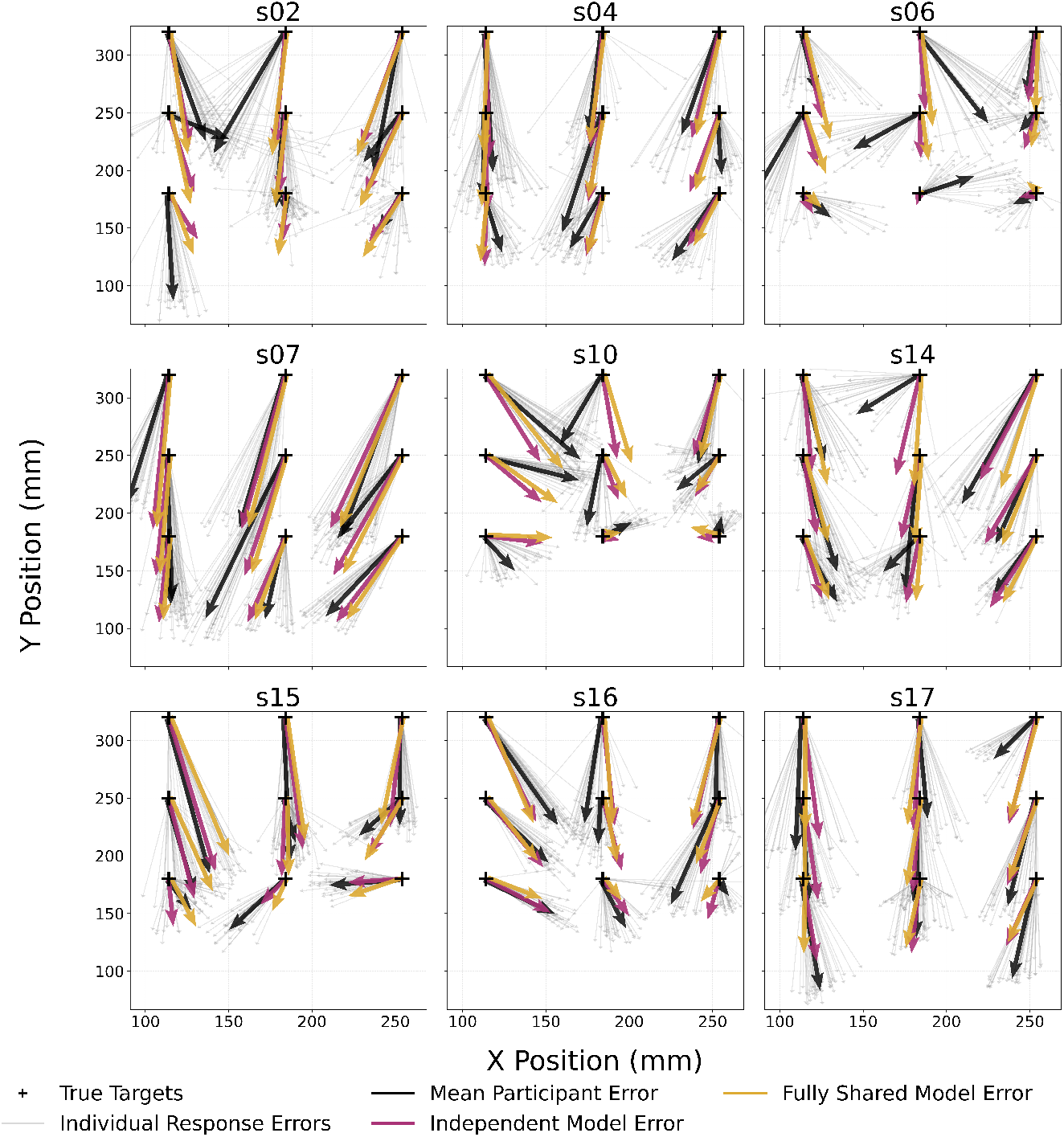
Tactile localization errors: Comparison of mean participant responses (black arrows) with Independent (red arrows) and Fully Shared Model (gold arrows) predictions across nine participants. For visualization clarity, responses from index and ring fingers have been spatially shifted and collapsed onto the middle finger position, allowing all 27 target locations and corresponding responses to be displayed in a 3×3 grid. The grid positions correspond to the 9 middle finger locations used in the experiment.

**Figure 6.**
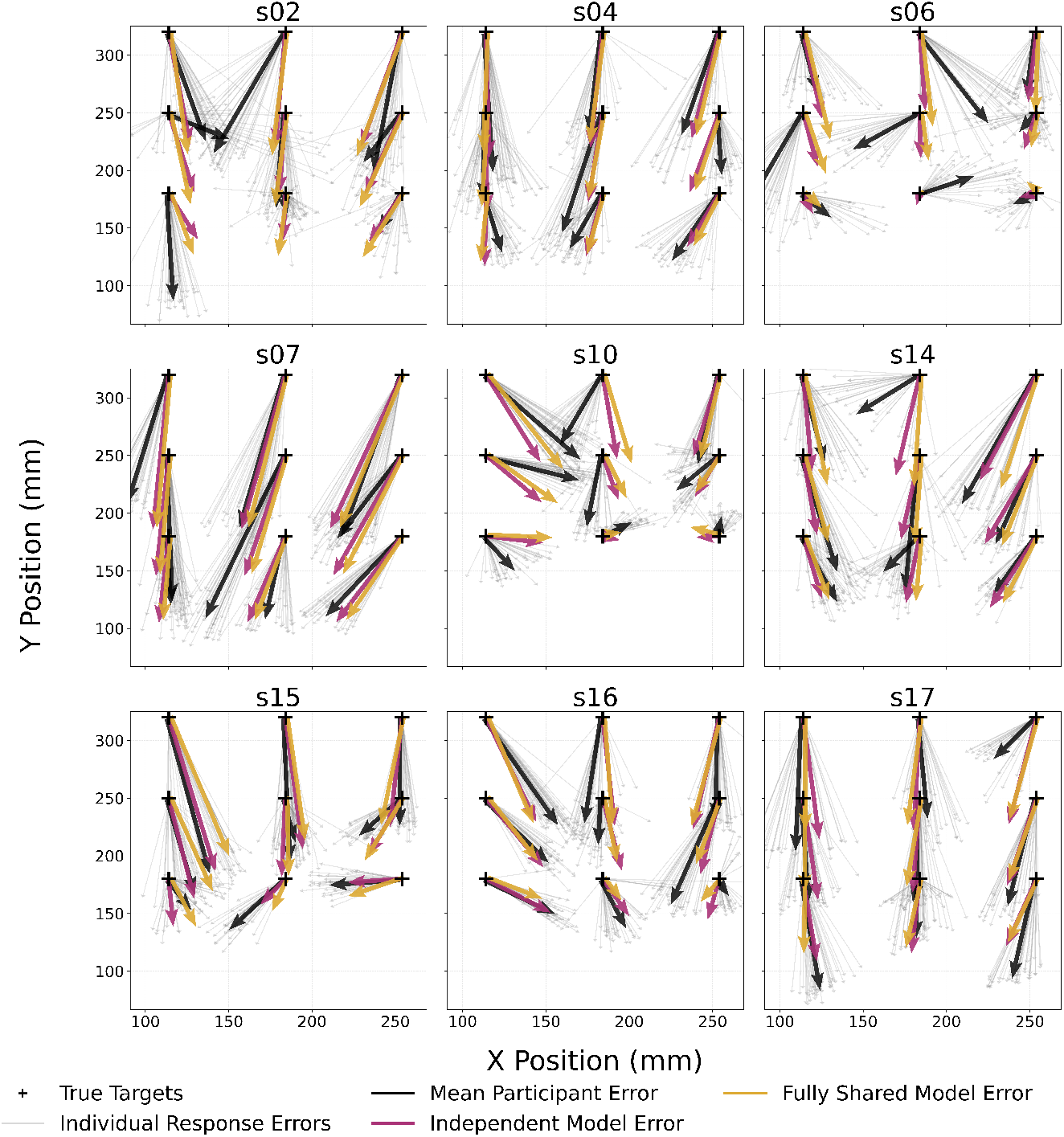
Proprioceptive localization errors: Comparison of mean participant responses (black arrows) with Independent (red arrows) and Fully Shared Model (gold arrows) predictions across nine participants. Data visualization is identical to Fig 5.

**Figure 7.**
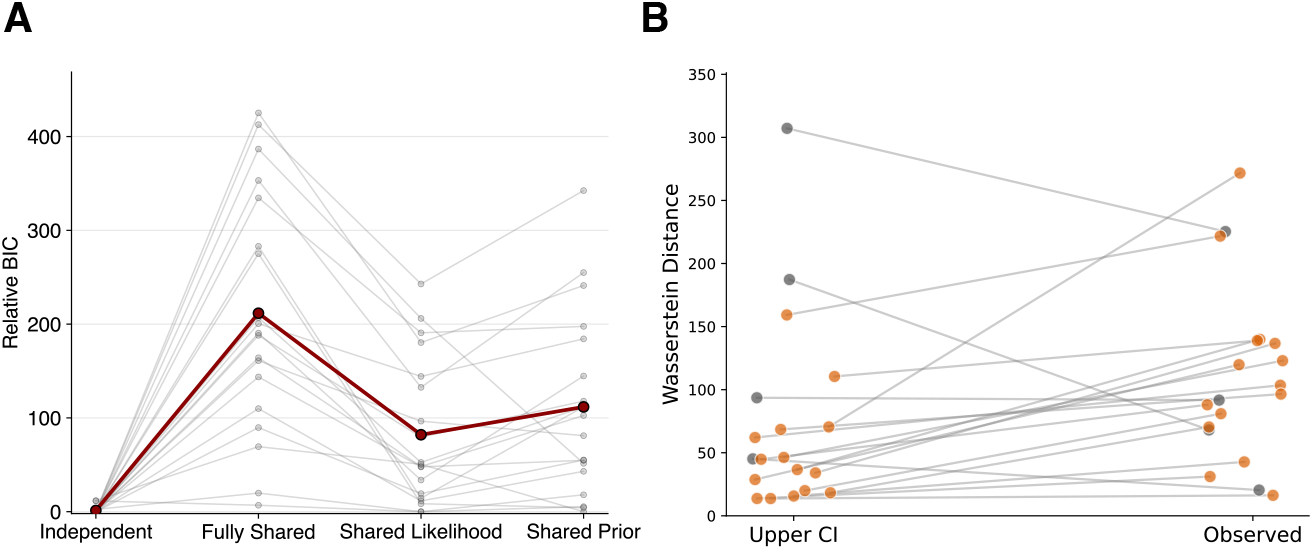
A) Model comparison across participants using Δ*BIC* relative to the best model for each participant. Each grey dot represents an individual participant’s Δ*BIC* for a given model, and the red dots represent the mean. B) Observed Wasserstein-2 distances versus the upper 95th percentile thresholds from null distributions for all participants (1000 shuffles per participant). Orange points dots indicate participants with significantly different priors between modalities, whereas grey dots indicate participants with non-significant differences

The Independent Model also outperformed the Shared Prior Model in 17 of 19 participants (mean Δ*BIC* : −110; range: [-342, 11]; mean BF: 9.19 ×10^23^), while the Shared Likelihood Model outperformed the Shared Prior Model in 15 of 19 participants (mean Δ*BIC* : −29; range: [-122, 154]; mean BF: 2.73 × 10^6^). Among integration-supporting models, the Independent Model was superior to the Shared Likelihood Model in 17 of 19 participants (mean Δ*BIC* : −801; range: [-242, 11]; mean BF: 3.37 ×10^17^), favoring models with greater parameter independence between modalities.

Finally, we assessed the Independent Model’s overall explanatory power using multiple pseudo-*R*^2^ measures relative to a null model. The Independent Model achieved moderate explanatory power across participants (McFadden’s pseudo-*R*^2^: 0.15 ± 0.03; Cox-Snell pseudo-*R*^2^: 0.43 ± 0.09).

A crucial distinction between the cueing and integration hypothesis is the nature of the prior expectations underlying tactile and proprioceptive localization. That is, the cueing hypothesis posits that priors are modality-independent, whereas the integration hypothesis suggests that they are modality specific. While the Δ*BIC* comparison suggested that the Independent Model provided better fit for most participants, this could also result from improved model fit estimation without substantially different priors. Fig 8 shows the modality-specific prior distributions over the workspace of example participants. These plots suggest differences in the spatial biases captured by the Independent Model for tactile and proprioceptive localization. Going beyond visual inspection, we therefore sought to quantitatively confirm that fits from the Independent Model captured statistically significant modality-specific priors, as this would be evidence against the cueing hypothesis. To examine whether the fitted spatial priors differed beyond what might be expected from random variation, we quantified the magnitude of differences between modality-specific priors using the Wasserstein-2 metric with permutation testing. Permutation testing of the distances between the modality-specific priors showed that 15 of 19 participants (78.9%) exhibited significant differences between tactile and proprioceptive prior distributions (p < 0.05; Fig 7B), suggesting that observed modality-specific spatial representations cannot be attributed to random variation. This clearly demonstrates that the fitted modality-specific prior distributions cannot be attributed to random variation.

**Figure 8.**
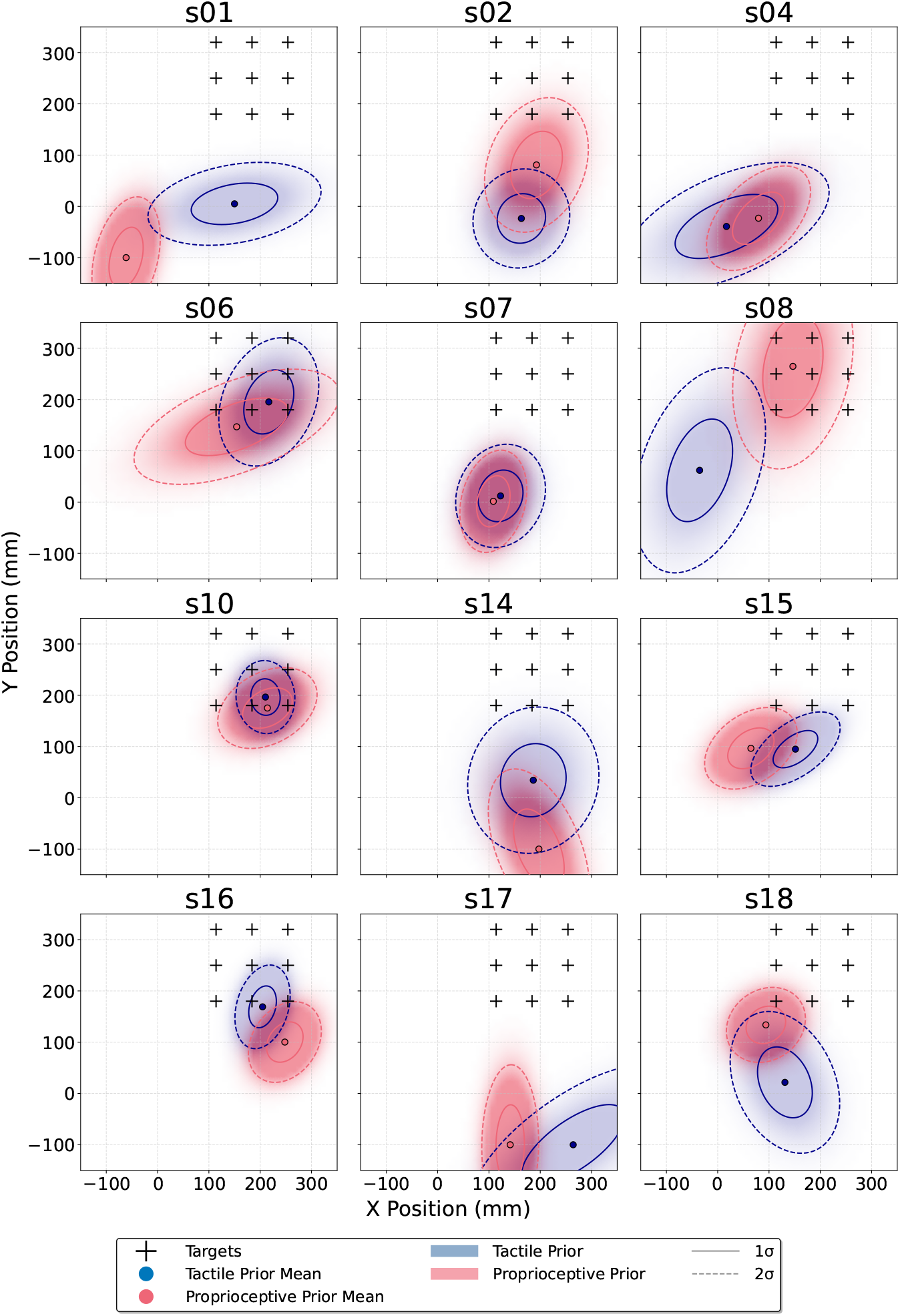
Fitted spatial priors from the Independent Model for representative participants. Each panel depicts one participant’s workspace where black crosses mark target locations. Colored regions represent 2D Gaussian prior distributions with their means: red shading illustrates the covariance ellipse of the tactile priors; blue shading illustrates the covariance ellipse of the proprioceptive priors.

## Discussion

In this study, we combined psychophysics (tactile and proprioceptive localization in 2D space) and Bayesian modeling to determine the computational underpinnings of tactile localization. This approach allowed us to adjudicate between two competing hypotheses: the integration hypothesis [2], which proposes that tactile inputs are remapped into external spatial coordinates via integration with proprioception; and the cueing hypothesis [23], which suggests that touch merely indicates a body location that is then localized through proprioceptive mechanisms—in this case, no remapping occurs. Both hypotheses make crucial distinctions between the structure of Bayesian computations underlying localizing both somatosensory modalities. Specifically, the integration hypothesis allows for distinct spatial priors, whereas the cueing hypothesis requires identical priors for tactile and proprioceptive localization. To differentiate between the two hypotheses, we implemented and compared four Bayesian model variants with different parameter-sharing structures, from fully independent parameters (supporting the strong integration hypothesis) to fully shared parameters (supporting the strong cueing hypothesis).

We found significant evidence in favor of the integration hypothesis, demonstrating that tactile and proprioceptive localization rely on distinct computational mechanisms. Indeed, the best-fitting model was the Independent Model, which was the most extreme version of the integration hypothesis. Crucially, most participants showed significantly different spatial prior distributions between tactile and proprioceptive localization, providing direct computational evidence against the cueing hypothesis. Our results challenge the view that tactile localization is computationally equivalent to proprioceptive localization [21], and instead suggest that tactile localization involves multimodal integration that remaps tactile inputs into external spatial coordinates.

More broadly, these results suggest that somatosensory spatial localization involves multiple, specialized representational systems that maintain distinct computations, rather than relying on a single shared spatial framework.

An important distinction of our approach from other studies [3, 5, 25, 26, 23] was the use of Bayesian modeling to test competing hypotheses about tactile and proprioceptive localization mechanisms. While Bayesian modeling has been successfully applied to various aspects of sensory processing [35, 33, 51, 52, 53], its application to body representation and tactile-proprioceptive integration remains sparse [38, 39, 42]. By decomposing observed behavior into its computational components [31, 30], we were able to gain insight into the spatial architecture of touch and proprioception that would not have been possible otherwise. Our study therefore highlights the theoretical benefits of using modeling to distinguish between competing hypotheses.

We observed considerable differences between the fitted priors for tactile and proprioceptive localization (Fig 7B, Fig 8). It is therefore important to consider possible computational and contextual factors that may contribute to these differences. Our results suggest that tactile localization requires a different series of coordinate transformations than proprioceptive localization. While proprioceptive localization involves transforming joint angles and limb configurations directly into external spatial coordinates [41], tactile localization might first map touch from skin-based coordinates to limb-based representations (Miller et al., 2022)—only then is it possible to integrate touch with proprioceptive information to achieve external spatial localization [1, 2, 7, 54]. Each additional transformation step likely introduces systematic errors that accumulate, potentially leading to the distinct spatial biases that manifest as different prior distributions [55, 56, 57].

Differences in environmental statistics could provide a contextual basis for distinct priors across different modalities, including vestibular [58], audition [59], and vision [60, 61]. Given that tactile and proprioceptive localization often occur in different spatial contexts, it is reasonable to assume that they may also have distinct statistics. When tactile events occur on the fingertips, the hands are often positioned to interact with objects or explore surfaces. These functional positions may differ from typical resting arm configurations [43]. These differences in environmental statistics could favor distinct spatial priors. However, maintaining separate priors could entail computational costs in neural resources. The existence of modality-specific priors in our data suggests these costs may be outweighed by functional benefits for tactile localization.

Localizing a touch in external space involves several multimodal regions throughout frontal and parietal cortices [18, 20, 62]. The posterior parietal cortex (PPC) appears to be especially important in this ability [63] and has traditionally been thought to implement the remapping of tactile stimuli from skin-based coordinates into a spatiotopic representation [12, 16, 13, 14, 64]. Proponents of the cueing hypothesis would interpret these findings differently. Rather than a spatial representation of touch, activity within these regions reflects the coding of a specific location on the body.

Indeed, localizing proprioceptive targets involves similar regions of PPC [65]. Though our study was only behavioral, our findings do have implications for how to interpret neural data. Specifically, given that our study provides evidence for tactile remapping, at least some frontal and parietal regions are involved in tactile localization above and beyond localizing the body.

This further raises the question of how the elements of our model might be implemented in the brain. We found that tactile and proprioceptive localization involved distinct sources of sensory evidence. This is consistent with what we know about the earliest sensory processing of both modalities. Touch and proprioception are only integrated once signals have reached Areas 1 and 2 of primary somatosensory cortex [1], at least 70 ms after sensory input [66]. Furthermore, our Bayesian modeling revealed that most participants exhibited significantly different spatial priors between tactile and proprioceptive conditions. These modality-specific spatial priors might be encoded as systematic differences in the preferred spatial locations of neural populations, with different tuning curve centers [67, 68] or response patterns between tactile and proprioceptive networks [69, 70]. The encoding of such priors is likely reflected in how the responses of neural populations vary across space. Recent findings show that tactile and proprioceptive space is represented by spatial gradients in overall response magnitude [71, 72, 73]. Priors could be implemented in biases of these gradients in the direction of spatial expectations.

While our results support the integration hypothesis over the cueing hypothesis, several limitations should be acknowledged. First, our modeling framework examined the outcomes of remapping processes in Cartesian space rather than the transformations themselves. We did not measure joint angles or body segment lengths, which would have enabled direct modeling of the kinematic transformations from proprioceptive signals to spatial coordinates [41]. Furthermore, our Bayesian models assumed Gaussian distributions for both priors and likelihoods. While this assumption enabled tractable modeling, real neural representations likely involve more complex, non-Gaussian distributions, (for instance due to signal dependent noise in joint angles [74]). Future research should address these limitations through more naturalistic experimental paradigms utilizing virtual reality or multisensory environments, and computational modeling that explicitly characterizes the full kinematic chain from sensory input to spatial output.

To conclude, we provided computational evidence that tactile and proprioceptive localization rely on distinct spatial representations. By directly comparing each mode of localization, we were able to adjudicate between two competing hypotheses of how touch is localized in external space. Indeed, Bayesian modeling demonstrated that tactile and proprioceptive localization relied on distinct spatial expectations about stimulus location. This finding strongly suggests that localizing a touch on the body and localizing the body itself are not the same process, a conclusion that has important implications for how we understand these everyday behaviors and their neural mechanisms.

## Acknowledgments

LEM and HOE are supported by ERC Starting Grant (101076991 SOMATOGPS). WPM is supported by NWA-ORC-1292.19.298, NWO-SGW-406.21.GO.009 and Interreg NWE-RE:HOME.

